# Gene networks and expression quantitative trait loci associated with platinum-based chemotherapy response in high-grade serous ovarian cancer

**DOI:** 10.1101/740696

**Authors:** Jihoon Choi, Anastasiya Tarnouskaya, Sean Nesdoly, Danai G. Topouza, Madhuri Koti, Qing Ling Duan

## Abstract

**Background:** A major impediment in the treatment of ovarian cancer is the relapse of platinum-resistant tumors, which occurs in approximately 25% of patients. A better understanding of the biological mechanisms underlying platinum-based response will improve treatment efficacy through genetic testing and novel therapies.

**Methods:** Using data from high-grade serous ovarian carcinoma (HGSOC) patients in the Cancer Genome Atlas (TCGA), we classified those who remained progression-free for 12 months following platinum-based chemotherapy as “chemo-sensitive” (N=160) and those who had recurrence within six months as “chemo-resistant” (N=110). Univariate and multivariate analysis of expression microarrays identified differentially expressed genes and co-expression gene networks associated with chemotherapy response. Moreover, we integrated genomics data to determine expression quantitative trait loci (eQTL).

**Results:** Differential expression of the Valosin-containing protein (VCP) gene and five co-expression gene networks were associated with chemotherapy response in HGSOC. VCP and the gene networks contribute to protein processing in the endoplasmic reticulum, which has been implicated in chemotherapy response. These findings were successfully replicated using independent replication cohort. Furthermore, 192 QTLs were associated with these gene networks and *BRCA2* expression.

**Conclusion:** This study implicates both known and novel genes as well as biological networks underlying response to platinum-based chemotherapy among HGSOC patients.

## Introduction

Ovarian cancer is the most lethal gynecological malignancy and the 8^th^ leading cause of cancer death in women around the world.^1^ According to the Global Cancer Observatory report in 2012, ovarian cancer accounts for 3.6% of all cancer cases and 4.3% of all cancer related deaths worldwide.^2^ High-grade serous ovarian carcinoma (HGSOC) is the most malignant form of ovarian cancer that accounts for up to 70% of all cases.^3^ Routine diagnosis is often difficult due to the lack of mass screening methods and heterogenous manifestations of the cancer symptoms, which result in approximately 75% of patients diagnosed with advanced stages.^4^ The average 5-year survival rates are 39% for Stage 3 and 17% for Stage 4 cancers.^1^

The current standard of care for ovarian cancer is aggressive cytoreductive surgery followed by platinum-based chemotherapy.^4^ However, this standard of care is not effective for all patients, with approximately 25% experiencing relapse within six months following platinum-based therapy, likely due to the development of antineoplastic resistance.^5^ The median survival time for recurrent ovarian cancers range from 12-24 months.^6,7^ Treatment options for patients with recurrent ovarian cancer include non-platinum-based chemotherapy regimens, immunotherapy, and molecular targeted therapy.^7,8^

Ovarian cancer has a multifactorial etiology that includes genetic and non-genetic risk factors. An estimated 23% of cases are hereditary, but the majority are sporadic with multiple reported risk factors such as history of gravidity, infertility, and late age menopause.^9,10^ A better understanding of the etiology of ovarian cancer, as well as the genetic mechanisms underlying variable response to platinum-based chemotherapy is needed for improved diagnosis and treatment. For example, previous studies reported that *BRCA1* and *BRCA2* genes, associated with increased risk of ovarian cancer, harbor mutations associated with platinum drug sensitivity and survival time.^11^ Similarly, tumor suppressor genes such as *RB1, NF1, RAD51B, PTEN* have been associated with acquired chemotherapy resistance.^12^ Earlier studies have also highlighted the importance of the immune system in the treatment of ovarian cancer. For example, loss of chemokines and disruptions to the IFN-γ pathway have been associated with poor treatment outcomes in HGSOC paients^13^ whereas the NFκB signaling pathway and elevated expression of *STAT1* were associated with increased response to platinum therapy.^14,15,16^ However, these known genetic variations do not account for all of the variability in chemotherapy response among HGSOC patients and there is currently no screening method to accurately predict prognosis prior to start of chemotherapy. Thus, further studies are necessary to determine additional modulators of chemotherapy response, which can be used as biomarkers for genetic testing.

The majority of earlier studies of chemotherapy response in ovarian cancer patients used univariate analysis of gene expression data known as differential gene expression (DGE) analysis.^17,18^ This analysis assumes that each gene functions in isolation within the genome and fails to capture the effects of complex gene-gene interactions. This study applies a multivariate approach to identify groups of interconnected genes, which may contribute to common biological pathways. These genes may each have modest effects that are not detected by conventional univariate analysis. Specifically, we applied Weighted Gene Co-expression Network Analysis^19^ (WGCNA), which uses an unsupervised machine-learning algorithm to identify clusters of highly correlated or co-expressed genes. Moreover, we correlated sequence variations with co-expressed gene networks to identify expression Quantitative Trait Loci (eQTLs), which are potentially regulatory variants associated with co-expression gene networks. A better understanding of the biological mechanisms regulating chemotherapy response will enable more effective treatments by improving the accuracy of genetic testing and through the identification of novel therapies for cancer patients.

## Methods

### Patient Classification

From the Cancer Genome Atlas (TCGA) Genomic Data Commons (GDC) portal^20^, we retrieved 587 high-grade serous ovarian carcinoma (HGSOC) patients with available clinical data using the TCGAbiolinks R/Bioconductor package.^21^ The interval between patient’s last primary platinum treatment and the onset of a recurrent tumor or progression of an existing tumor was used as a metric for determining chemotherapy sensitivity. Patients who developed a new tumor in less than six months following their last platinum treatment were defined as resistant (N=110). In contrast, those who did not have a recurrent tumor event for over a year after their last primary platinum treatment were defined as sensitive (N=160). Individuals who had a recurrent tumor event between six months to one year following chemotherapy were excluded from the study. This strategy for dichotomizing resistant from sensitive patients was used to enrich for the genetic differences.

### Transcriptomics Data Processing and Analysis

#### Expression Microarrays

Of the 270 HGSOC subjects classified as sensitive or resistant to chemotherapy, 238 (138 sensitive, 100 resistant) had microarray expression data available (Affymetrix ht_hg_u133a chip) in the GDC portal. The robust multi-array average (RMA) method^22^ in the *affy* package from Bioconductor^23^ was used for background correction, log-transformation, and quantile normalization of the probe intensities. Two potential outliers and two duplicated samples were removed from the study during the quality control step using the *arrayQualityMetrics*^24^ package (see **Supplemental Data 1** for steps of pre-processing), resulting in 135 sensitive and 99 resistant HGSOC subjects in the expression set. Next, probes were filtered using the median absolute deviation (MAD) whereby the top 50% with highest variation (n=11,107) were selected for analysis. Thus, this non-specific filtering step removed probes with no or low variability, which are not likely to be differentially expressed between sensitive and resistant patients. Moreover, this step reduces the number of multiple testing corrections and therefore, the likelihood of false positives.

#### Differential Gene Expression Analysis

The *Limma*^25^ package in Bioconductor^26^ was used to identify differentially expressed genes between chemo-sensitive and resistant groups using linear models. The false discovery rate (FDR) method was employed as a measure for multiple testing correction to control for type I error. Age at diagnosis and stage of tumor were included as covariates in this analysis (**Supplemental Data 2**).

#### Weighed Gene Co-expression Network Analysis (WGCNA)

We performed hierarchical clustering of genes using the R package *WGCNA*^19^, which groups genes based on their similarity in expression. This was achieved by first creating a similarity matrix using Pearson correlations of expression among all genes. The resulting matrix was raised to a power of 9, as suggested by the soft-thresholding power estimation plot (**Supplemental Figure 1**). This assumes that raising the correlation matrix to a power will enrich differences between weak and strong signals, allowing for better quantification of gene-gene interactions. In addition, because information between two genes are often too weak and noisy to interpret, similarity matrix is transformed to Topological Overlap Matrix (TOM), where strength of association between a pair of genes is reinforced by the common neighbors shared by them. To avoid excessive splitting of genes into smaller modules, minimum module size was set to 30, split sensitivity (deep split) to 4, and modules with similar expression profiles were merged at a height of 0.5 (**Supplemental Figure 2**). Using principal component analysis, we calculated the module eigengene for each co-expression cluster to summarize module expression into a single measure. Each module eigengene was tested for association with platinum chemotherapy response using generalized linear models, while adjusting for age at diagnosis and stage of cancer as covariates. Finally, we used *Cytoscape*^27^, an open source bioinformatics platform, to visualize gene co-expression networks.

#### Gene function and pathway annotations

The Database for Annotation, Visualization and Integrated Discovery (DAVID) ^28^ was applied to identify biological pathways and functions that were enriched in each significant gene co-expression module. We also screened significant genes in the GeneMANIA^29^ database to identify for functional connections reported in published literature. Next, we searched the UCSC TFBS conservation sites track with DAVID to identify enriched motifs of transcription factors that may co-regulate genes within each cluster. Finally, we used Drug–Gene Interaction database (DGIdb)^30^, a public database with curation of data describing relationships between genes, chemicals, drugs, and pathological phenotype, to identify genes with prior reported associations with chemotherapeutic agents.

#### Validation of differentially expressed gene

The Kaplan–Meier plotter was used to cross-validate the effect of differentially expressed genes in an independent ovarian cancer cohort (Geo accession identifier: GSE9891).^31^ This replication cohort included gene expression profiling (Affymetrix Human Genome U133 Plus 2.0 Array) of 285 ovarian tumor samples. Patients were filtered to include those with cancer histology of serous carcinoma and who received chemotherapy containing a platin compound to allow close comparison with the TCGA ovarian cancer cohort. This step omitted a total of 60 subjects from analysis, which included 21 with endometrioid carcinoma cases and 43 who did not receive platinum therapy (4 overlapping subjects). Thus, 225 patients remained for replication analysis. Patient survival was evaluated using Cox proportional hazards model and progression-free survival (PFS) was the primary outcome used in the replication analysis.^32^

#### Validation of co-expression network

For validation of co-expression networks, SurvExpress database was used, which allow users to validate the combined effect of multiple gene expression measures with a target trait.^33^ Same cohort and filtering steps were used for validation (GSE9891, N = 225). Survival curve was evaluated using Cox proportional hazards model and progression-free survival (PFS) was the primary outcome used in the replication analysis.

### Genomics Data Processing and Analysis

#### Genomics Data

Single nucleotide polymorphisms (SNPs) data from the germline tissues (DNA extracted from blood or solid non-tumor ovarian tissue) were obtained from the TCGA legacy database. The Affymetrix Genome-Wide Human SNP Array 6.0 was used to capture genetic variations, which detected 906,600 SNPs. Of the 270 subjects from TCGA classified as resistant or sensitive to platinum-based chemotherapy, 266 (157 sensitive and 109 resistant) had genotype data available.

#### Imputations

The imputation of autosomal chromosomes was performed using the Michigan imputation server pipeline^34^. We used the 1000 Genome Project phase 3 sequencing data (version 5)^35^ reference panel for the imputation of missing genotypes. We then used Eagle v.2.3^36^ for phasing of the genotypes to their respective chromosomes. For the imputation of variants on the X chromosome, SHAPEIT^37^ was used for phasing in combination with the 1000 genomes project phase 3 (version 5) reference panel (**Supplemental Data 3**).

#### Quality control

##### Subject level

Two pairs of individuals had a relatedness coefficient (pi-hat) > 0.9, which are likely duplicated samples. One subject from each pair was randomly removed from the dataset. Next, inbreeding coefficients (F) were computed for each subject using PLINK^38^. A total of 18 subjects with high homozygosity (F>0.05) or heterozygosity (F<-0.05) rates were excluded. Moreover, genetic sex was estimated based on heterozygosity rates (F) of the X chromosome, and four subjects who had undefined (F>0.2) genetic sex were removed from the study.

##### SNP level

SNPs with minor allele frequencies (MAF) less than 1% or with genotyping call rate less than 90% were removed. This step removed 38,430,595 SNPs with MAF < 0.01, resulting in 9,528,963 SNPs to be used for further analysis.

#### Genome-wide Association Study

After imputations and quality control, 240 subjects (N= 142 sensitive, 98 resistant) and a total of 9,528,963 SNPs (MAF > 0.1) remained available for analysis (**Supplemental Data 4**). Plink (v.1.90) was used to compute genome wide and BRCA1/2 targeted association analysis using a logistic regression model. Age at diagnosis and tumor stages were included as covariates.

### Expression Quantitative Trait Loci (eQTL) Analysis

Common SNPs (MAF > 0.01) were tested for association with differentially expressed genes and gene networks using the *matrixeQTL* R package^39^. Loci correlated with nearby gene expression indicates potential regulatory function of the SNP on the corresponding gene, known as cis-expression Quantitative Trait Loci (cis-eQTL). Cis-eQTLs are defined as correlated SNPs found within 1 Mb from the gene transcriptional start site (TSS).

## Results

Differential gene expression (DGE) analysis identified that low expression of a probe (208648_at) mapping to the Valosin Containing Protein (*VCP*) gene was significantly associated with resistance to chemotherapy (FDR adjusted p-value < 0.05; **Figure 1**). In addition, 606 probes mapping to 521 unique genes were nominally correlated with chemotherapy response (unadjusted p-value < 0.05; **Supplemental Table 1**). The Kaplan– Meier survival plotter demonstrated that low expression of VCP is associated with poor progression-free survival (p = 0.015) and shorter median survival time (**Figure 2**) in an independent ovarian serous cancer cohort following treatment with platinum antineoplastic agents. Functional annotation analysis showed that the 521 genes were enriched for numerous oncogenic processes, such as cellular response to DNA damage stimulus (GO:0006974), DNA repair (GO:0006281), positive regulation of cell growth (GO:0030307), and positive regulation of apoptotic process (GO:0043065). Pathway analysis identified many immune related pathways including the mitogen-activated protein kinase (MAPK) and B-Cell antigen receptor (BCR) signaling pathways. A detailed list of functional annotations and pathways are shown in **Supplemental Data 5**.

**Figure 1.**
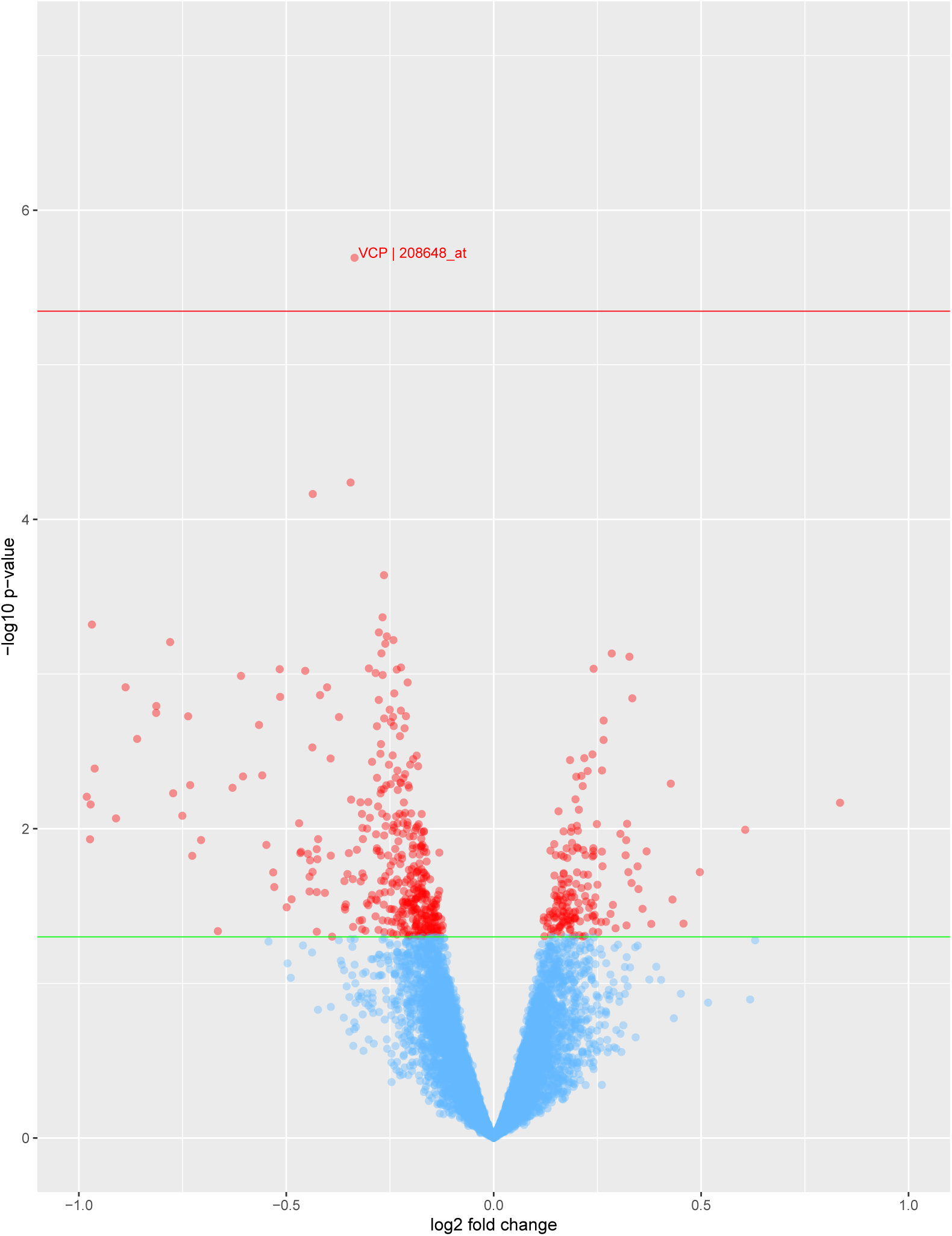
Differential expression analysis of platinum-based chemotherapy response in HGSOC patients. Volcano plot showing univariate association analysis results. One probe, 208648_at, which maps to the Valosin-Containing Protein (*VCP*) gene is significantly differentially expressed and correlated with chemotherapy outcome after multiple testing correction. Red horizontal line demonstrates FDR-corrected p value (< 0.05) threshold. A total of 606 probes mapping to 521 unique genes are also differentially expressed with nominal significance, as indicated by the green line (p = 0.05).

**Figure 2.**
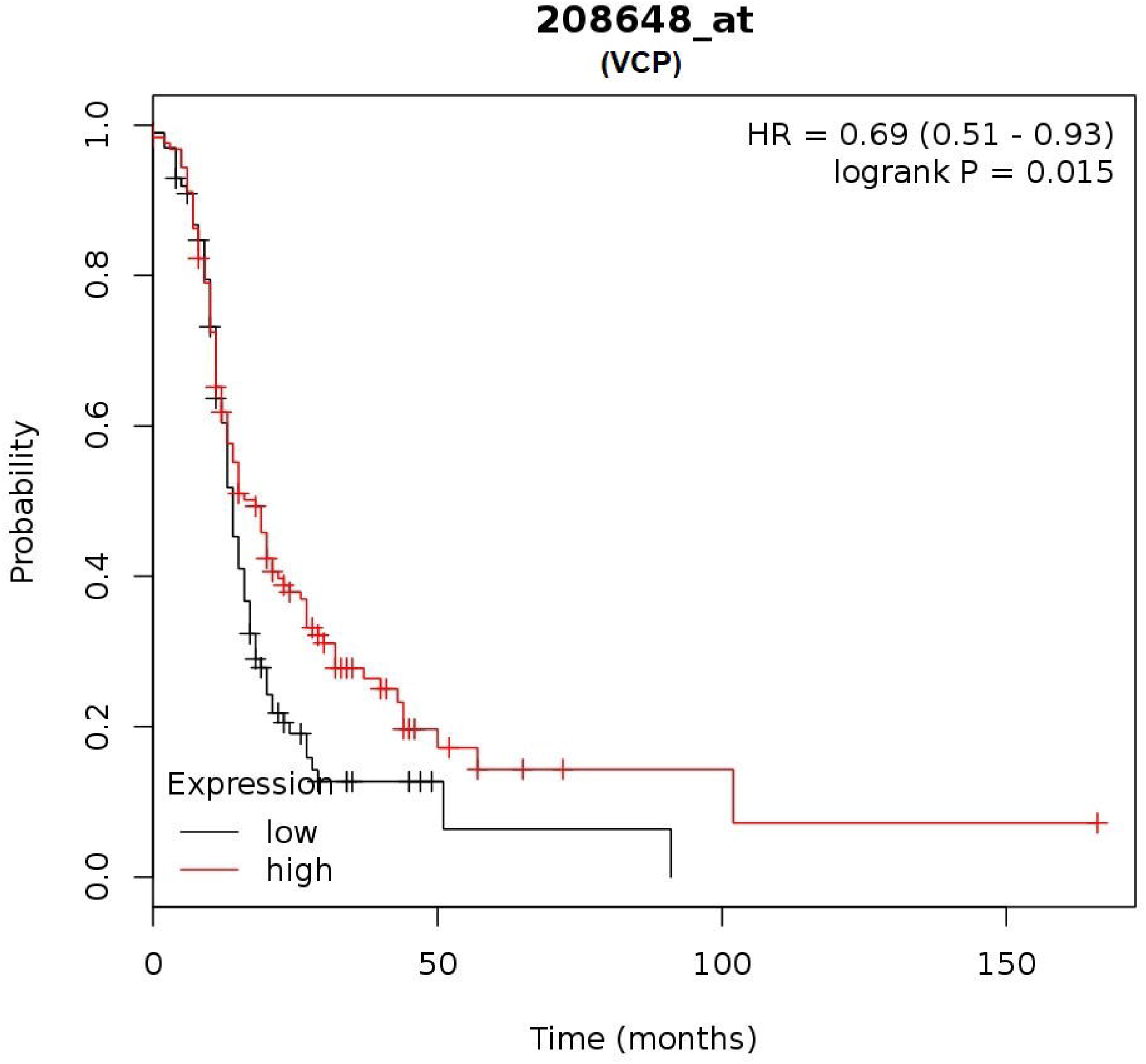
Kaplan- Meier plot of two ovarian cancer cohorts for cross-validation of differentially expressed gene VCP. Kaplan-Meier (KM) plot shows the Progression Free Survival (PFS) of the replication ovarian cancer cohort post platinum-based chemotherapy treatment. Red line indicates the PFS of patients with high VCP expression, while black line indicates the PFS of patients with low VCP expression. Patients with high expression of VCP is associated with better PFS with statistical significance.

The hierarchical clustering of genes using *WGCNA* resulted in 86 unique modules of co-expressed genes (**Supplemental Table 2**). Each module was assessed for association with chemotherapy response (results shown in **Supplemental Figure 3**). Five gene clusters (*honeydew1, lightcyan1, lightpink3, orangered4, and skyblue3*) were co-downregulated in platinum-resistant patients (p < 0.05); one gene cluster remained significant after Bonferroni correction (*honeydew1*, p-value=7e-05) (**Figure 3**). This signal was validated using SurvExpress in an independent cohort (Tothill et al.), which demonstrates that downregulation of the genes in five modules result in significantly reduced survival of patients (**Supplemental Figure 4**). The five significant modules were annotated using DAVID, which identified gene enrichment for biological pathways including apoptosis, negative regulation of the Wnt signaling pathway, transcription, immune responses, and DNA double-strand break processing involved in repair via single-strand annealing. GeneMANIA analysis showed that genes in these modules were previously reported in 49 publications, some of which documented associations with oncogenic pathways and chemotherapeutic outcomes. Our search of these genes in the gene-drug interaction database (DGIdb) found that 35 were associated with chemotherapeutic agents such as carboplatin and paclitaxel. Furthermore, we identified over-represented conserved transcription factor binding sites located in genes from each module. A detailed list of functional annotations, transcription factors and pathways related to gene modules can be found in **Supplemental Data 6.**

**Figure 3.**
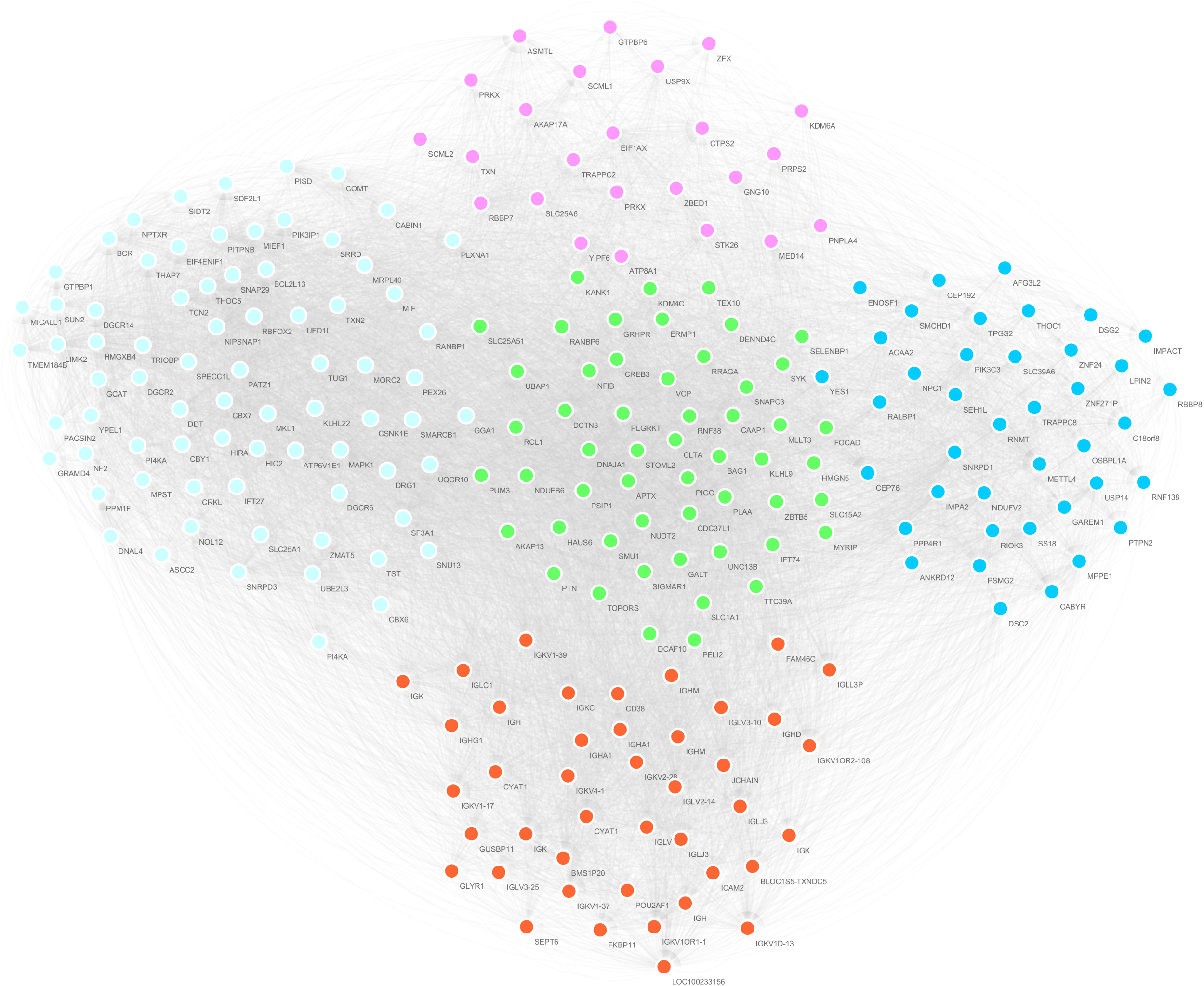
Gene co-expression modules correlated with platinum-based chemotherapy response. Network plot displaying the five significant gene co-expression modules from *WGCNA*: honeydew1 - centre, lightcyan1 - left, lightpink3 - top, orangered4 - bottom, and skyblue3 - right. Nodes represent probes and edges are connections among the probes. Co-expressed probes (i.e. belonging to a single module) are indicated by the same color.

Our GWAS of SNPs did not identify any variants correlated for platinum chemotherapy response after multiple testing correction. The Manhattan plot (**Supplemental Figure 5**), demonstrates that none of the SNPs meet the genome-wide significance threshold (p<5×10-8), as indicated in the by the red horizontal line. This is likely due to insufficient statistical power resulting from low number of subjects in TCGA-OV cohort. Next, we performed a targeted association analysis of two well-known genes associated with ovarian cancer and chemotherapeutic outcomes: *BRCA1* and *BRCA2*. Of the 238 SNPs in *BRCA1* and 256 in *BRCA2*, we identified 34 SNPs in *BRCA2* and 1 SNP in *BRCA1* that were significantly associated with chemotherapy response (**Supplemental Table 3**).

Next, SNPs were tested for correlation with expression of the 606 differentially expressed probes and 5 co-expression modules. This identified 5,242 cis-eQTL associated with the expression of differentially expressed probes, and 192 cis-eQTL associated with co-expression networks (**Supplemental Data 7**).

## Discussion

In this manuscript, we identified novel genes and gene networks correlated with variable response to platinum-based chemotherapy in HGSOC patients. Using a univariate analysis approach, we identified a differentially expressed locus encoding the valosin-containing protein (*VCP*) associated with sensitivity to platinum-based chemotherapy. In addition, we applied a multivariate, co-expression analysis method for identifying groups of interconnected genes that could contribute to common biological pathways. This method identified 5 clusters of co-expressed genes nominally correlated with chemo-response, one of which remained significant after correction of multiple testing. Genes in these modules have been annotated to be associated with biological pathways such as apoptosis, transcription, immune response, negative regulation of the Wnt signaling pathway and DNA double-strand break processing involved in repair via single-strand annealing.

The most significantly associated probe identified in the DGE analysis was for a gene encoding Valosin-containing protein (*VCP*, p = 3.91E-06). We have confirmed that this signal is replicated in an independent ovarian cancer cohort with statistical significance (p=0.015). *VCP* plays a critical role in disintegrating large polypeptide cellular structures for further degradation by proteolytic enzymes. It functions to regulate important pathways of DNA repair, replication and cell cycle progression by removing faulty polypeptide structures from chromatin material, ribosomes, endoplasmic reticulum and mitochondria. This gene has been previously identified as a potential biomarker for predicting the success of platinum-based chemotherapy in lung cancer patients.^40^

Many of the nominally correlated genes from our DGE analysis are enriched for pathways known to be critical for oncogenesis and chemotherapeutic drug resistance. For example, probe 213532_at mapping to the A Disintegrin and Metalloproteinase-17 (*ADAM17*) gene was upregulated in the chemo-resistant group (p = 0.017). This is consistent with prior studies, which reported that overexpression of *ADAM17* results in reduced cisplatin-induced apoptosis in hepatocellular carcinoma cells and may contribute to cisplatin resistance via the EGFR pathway^41^. Moreover, we identified another up-regulated probe 205239_at (p = 0.003) for the gene encoding Amphiregulin (AREG), a protein found in the EGFR signaling pathway, which has been reported to promote ovarian cancer stemness and drug resistance to anti-cancer therapy.^42^ Taken together, our findings support a potential role of the AREG-EGFR signaling pathway in the development of platinum-resistance in ovarian carcinoma.

It is also important to note that one of the most enriched pathways from our DGE analysis was the protein processing in endoplasmic reticulum (hsa04141) pathway. Using DAVID, we have identified 18 differentially expressed probes mapping to 15 unique genes within this pathway. Prior studies have highlighted that endoplasmic reticulum (ER) stress may cause cisplatin resistance in ovarian carcinoma by inducing autophagy in cancer cells, allowing them to escape apoptosis.^43,44^ Findings from this pathway analysis suggest that future studies of the contribution of ER stress in cisplatin sensitivity is needed to improve our understanding of platinum-resistance in ovarian cancer.

In our co-expression network analysis, the gene module “*honeydew1*” showed the most significant correlation with chemotherapy response (p = 6.53e-05). This association signal was validated with statistical significance (p = 5.88e-07) in an independent replication cohort. This module includes two probes that map to *VCP*, a gene that was associated with platinum-based chemotherapy response in our DGE analysis. Additional genes in this module were associated with positive regulation of mitochondrial membrane potential, protein ubiquitination, mitosis, alternative splicing, and apoptotic processes. Pathway analysis showed that this module is involved in protein processing in the endoplasmic reticulum. A prior study has found that *VCP* plays a crucial role in cancer cell survival through extraction and degradation of unfolded proteins in endoplasmic reticulum, and noted that lower expression of *VCP* was associated with poor response to platinum□based chemotherapy.^45^ In alignment with this finding, genes co-expressed in the *honeydew1* module were co-downregulated in chemo-resistant patients.

The *honeydew1* module is composed of 76 probes mapping to 54 unique genes, and of these, 44 genes are located in chromosome 9, demonstrating the importance of chromosome 9 in the regulation of chemo-resistance in ovarian cancer. These findings support previous studies, where genetic imbalance and alterations in chromosome 9 have been associated with progression of ovarian cancer and increased cisplatin resistance.^46^ Analysis of overrepresented transcription factor binding sites demonstrated that genes in this module may be co-regulated by a common transcription factor known as organic cation transporter 1 (OCT1). We found that over 96% of genes in this module (49/54 genes) contain a nucleotide motif bound by OCT1. Prior studies have reported that silencing OCT1 impaired cisplatin-induced apoptosis in esophageal cancer cells, and that cisplatin-resistant cells were already expressing significantly reduced levels of OCT1.^47^ Taken together, these findings characterize a network of co-expressed genes that is associated with platinum-resistance in ovarian cancer. Genes within this module may be co-regulated by the OCT1 transcription factor, which may be used as a novel potential target for ovarian cancer therapies

The other four co-expression modules, which we were also able to replicate in an independent cohort, include genes known to be involved in oncogenic process and drug response outcomes. For example, the *orangered4* module is composed of genes that are critical for regulation of immune response, which have been shown to play a role in chemotherapy response in HGSOC.^14,15^ Genes in this module are associated with functional annotation terms including immunoglobulin receptor binding, antigen binding, B cell receptor signaling pathway, and phagocytosis. In addition, 10 of the 58 genes in this module are enriched for a common transcription factor binding site: acute myeloid leukemia 1 protein (AML1). This transcription factor is involved in haematopoiesis process and immune functions such as thymic T-cell development. Studies have reported that the AML1 transcription factor is overexpressed in ovarian cancer patients, and may contribute to cancer cell proliferation, migration and invasion.^48^ In addition, we found that the *lightpink3* module is strongly associated with transcription regulation process, which plays a pivotal role in cancer progression. Finally, genes in modules *lightcyan1* and *skyblue3* are target regions of chemotherapeutic agents such as tyrosine kinase inhibitors (**Supplemental data 6**).

Targeted analysis of *BRCA1* and *BRCA2* SNPs demonstrated that 28 out of 35 variants associated with chemotherapy response were also cis-acting eQTLs, correlated with the expression of *BRCA2* as well as neighboring genes *N4BP2L1, N4BP2L2, FRY*, and *STARD13* (nominal p-value <0.05). Both *BRCA2* and *STARD13* are well known tumor-suppressors, and upregulation of *N4BP2L1* and *N4BP2L2* have been associated with positive prognosis in ovarian cancer cases.^49^ This finding shows the potential regulatory effect of the variants in *BRCA2*.

In addition, our annotation results show that 47% of platin-response associated variants in *BRCA2* are linked with LDL/HDL cholesterol levels (**Supplemental Table 3**). Prior studies of lung and ovarian cancers consistently reported that cholesterol levels may affect the efficacy of platinum-based chemotherapeutic agents.^50,51^ Our findings indicate a new link between genetic variants in *BRCA2* and platinum-based chemotherapy response through cholesterol level regulation.

## Conclusion

In this study, we identified genes and gene networks correlated with platinum-based chemotherapy response in high-grade serous ovarian cancer patients, which implicate known and novel biological mechanisms. Specifically, we identified that reduced expression of *VCP* is associated with platinum-resistance. This gene is critical for removing unfolded proteins from the endoplasmic reticulum and have been correlated with cancer cell survival and platinum-based chemotherapy response in prior studies. In addition, we report potentially regulatory variants in the *BRCA2* gene correlated with chemotherapy response and the expression of genes that determine cholesterol levels. Moreover, we identified a novel group of potentially co-regulated genes on chromosome 9 that are correlated with platinum resistance using a machine-learning algorithm. In addition, genes from this module are also involved in protein processing in the endoplasmic reticulum. This manuscript supports earlier studies which implicated *VCP* and *BRCA2* genes in chemotherapy response. We also identified regulatory variants in *BRCA2* and additional genes co-regulated with *VCP* on chromosome 9 that also contribute to protein removal in the ER. Findings from our study could facilitate genetic testing through the identification of gene signatures that may predict chemotherapy response as well as lead to novel drug targets, given a better understanding of the biological mechanisms underlying chemotherapy response.

## Acknowledgements

Computations in this manuscript were performed on resources and with support provided by the Centre for Advanced Computing (CAC) at Queen’s University in Kingston, Ontario. The CAC is funded by: the Canada Foundation for Innovation, the Government of Ontario, and Queen’s University. JC is funded by Queen’s University, Faculty of Health Sciences Dean’s Doctoral Award. QLD receives funding from the Canadian Institutes of Health Research and Queen’s University.

## Author contribution

J.C. performed the data analyses and drafted the manuscript. D.G.T., S.N. and A.T. assisted in the data analyses. Q.L.D. designed the research project, supervised data analyses and assisted in the writing of the manuscript. M.K. assisted in the study design and editing of the manuscript.

## Data availability

Transcriptomics, genomics, and clinical data used for the analysis of TCGA-OV cohort can be accessed/downloaded from the Genomic Data Commons (GDC) Data Portal (https://portal.gdc.cancer.gov/). Gene expression and clinical data of the replication cohort can be accessed/downloaded from Gene Expression Omnibus (GEO) database (https://www.ncbi.nlm.nih.gov/geo/query/acc.cgi?acc=GSE9899).

