## Supplemental Figure 1 for "Gene networks and expression quantitative trait loci associated with platinum-based chemotherapy response in high-grade serous ovarian cancer"

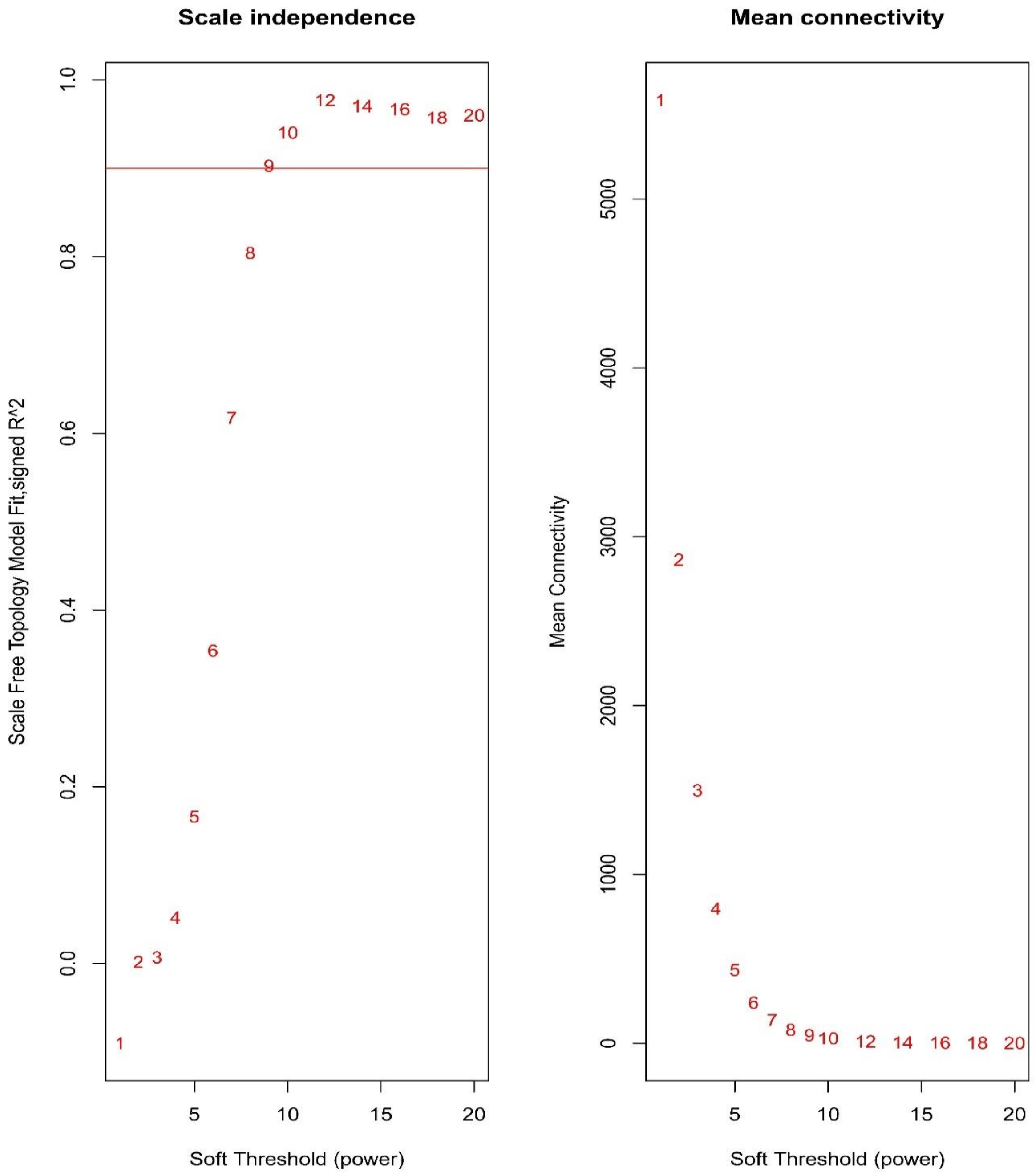


In this plot, we analyze the network topology at various soft thresholds (power). The assumption is that raising the similarity matrix by a power will enrich for differences between strong and weak signals. Scale independence plot (**left**) shows the change of scale-free fit index (r2) per change in power. The mean connectivity plot (**right**) shows the change of average connectivity between genes for each power increment. These results show that at power 9, scale free index reaches 0.9 (red horizontal line) and the network strongly resembles to a scale-free graph.
