## Supplemental Figure 2 for "Gene networks and expression quantitative trait loci associated with platinum-based chemotherapy response in high-grade serous ovarian cancer"

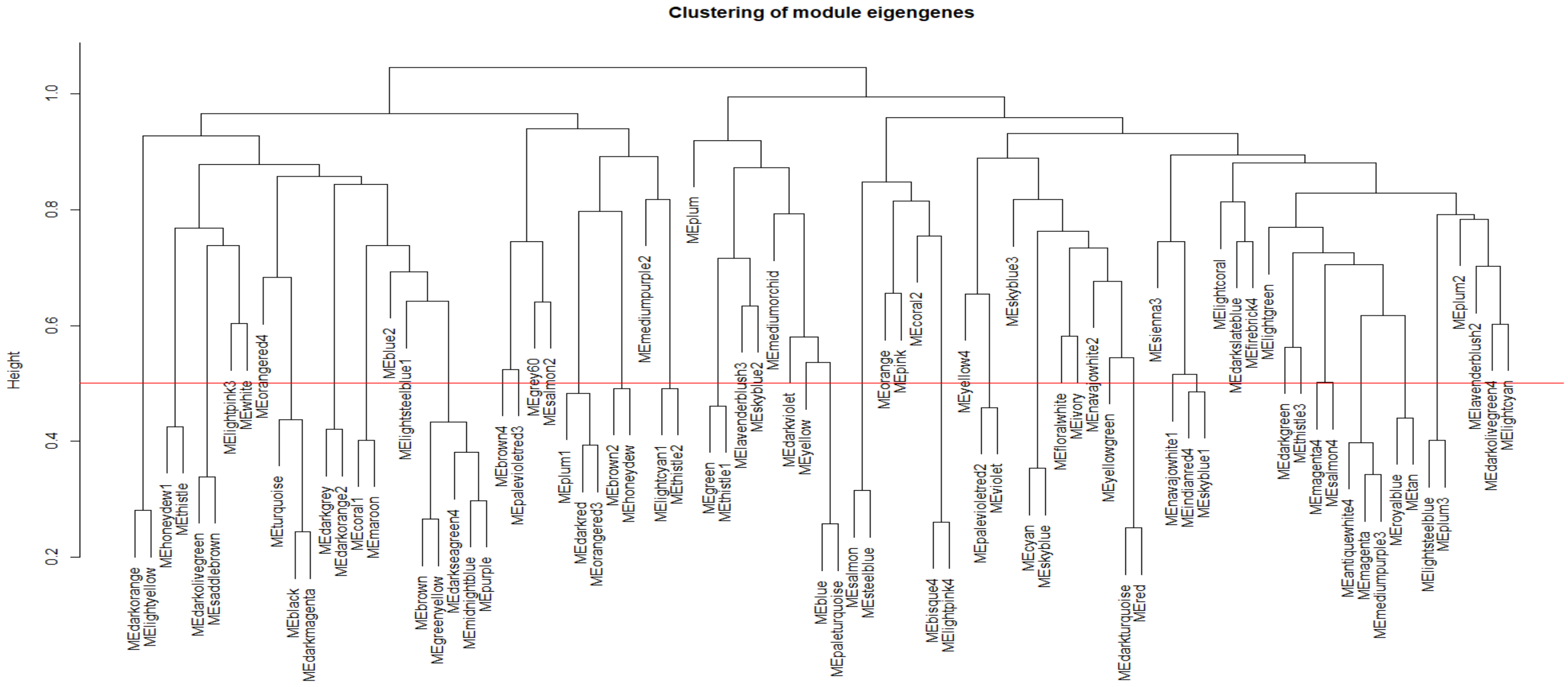


This plot shows every gene co-expression clusters identified from hierarchical clustering algorithm, where modules are represented by their eigengene. If similarity between two eigengenes are strongly correlated, we merge the two modules since genes in both modules would be strongly co-expressed. Red horizontal line in the plot indicates a height at which we merge modules.
