## Supplemental Figure 3 for "Gene networks and expression quantitative trait loci associated with platinum-based chemotherapy response in high-grade serous ovarian cancer"

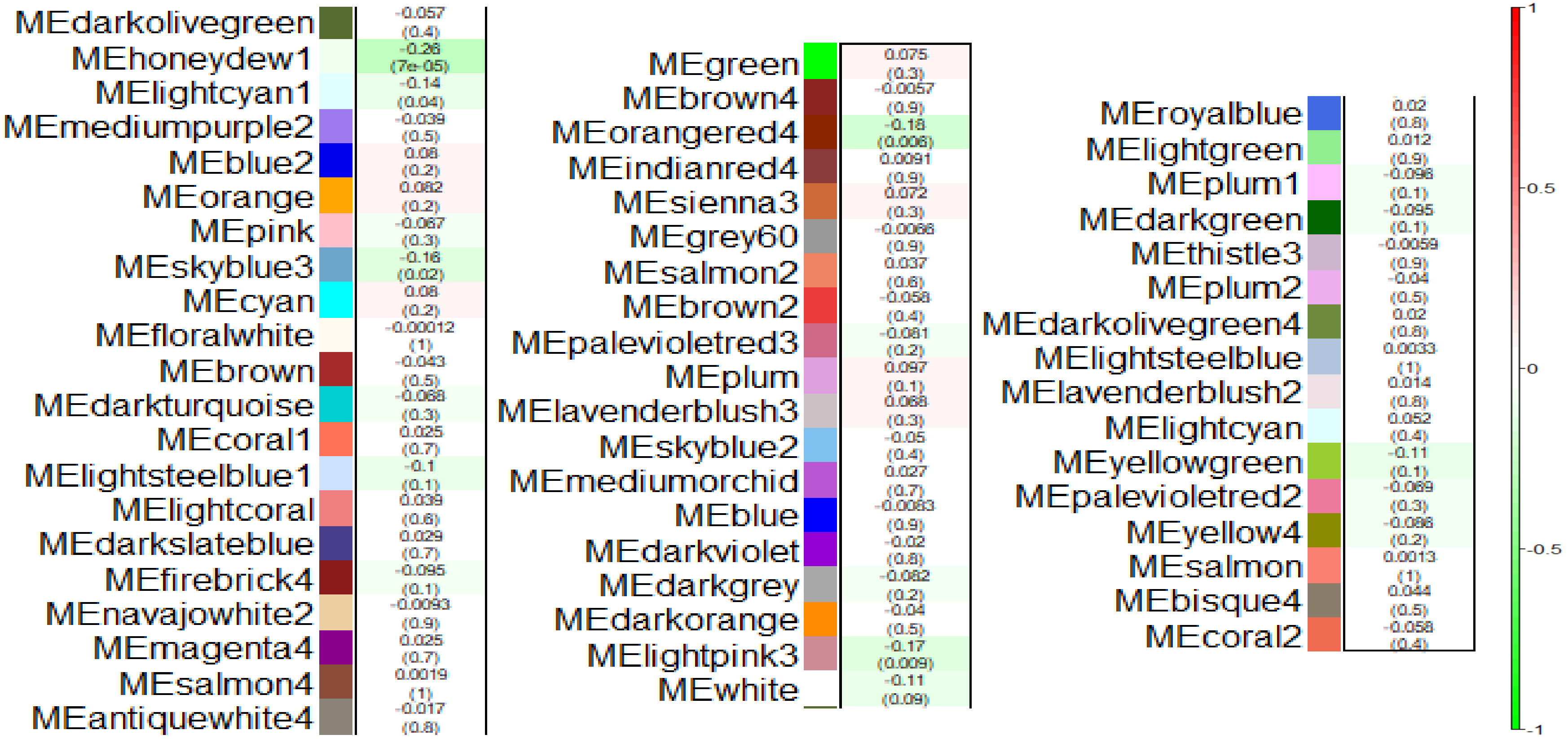


This plot shows heatmap of module-trait correlation analysis. Each module eigengene was tested for association with chemotherapy response using generalized linear model. Significance (p-value; in bracket) and strength of relationship (r2) between module and chemotherapy response is indicated in the plot. Red colored modules are expressed higher in resistant population, whereas green modules are expressed higher in sensitive population. Intensity of the color indicates strength of association and its statistical significance.
