## Supplemental Figure 4 for "Gene networks and expression quantitative trait loci associated with platinum-based chemotherapy response in high-grade serous ovarian cancer"

### Honeydew1 module replication (53 unique genes)

**Concordance Index = 70.5, Log-Rank Equal Curves p=1.368e-07, R^2=0.177/0.986**
**Risk Groups Hazard Ratio = 2.88 (conf. int. 1.9 ~ 4.36), p=5.881e-07**

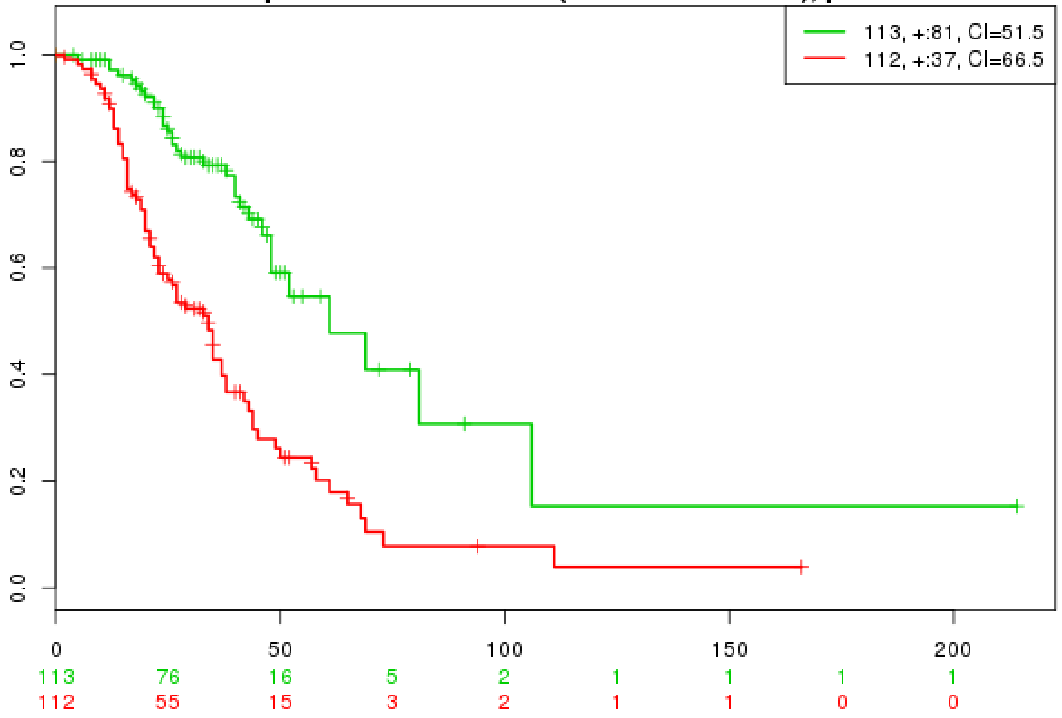


### LightCyan1 module replication (76 unique genes)

**Concordance Index = 75.94, Log-Rank Equal Curves p=7.938e-14, R^2=0.358/0.986**
**Risk Groups Hazard Ratio = 4.59 (conf. int. 2.96 ~ 7.1), p=9.04e-12**

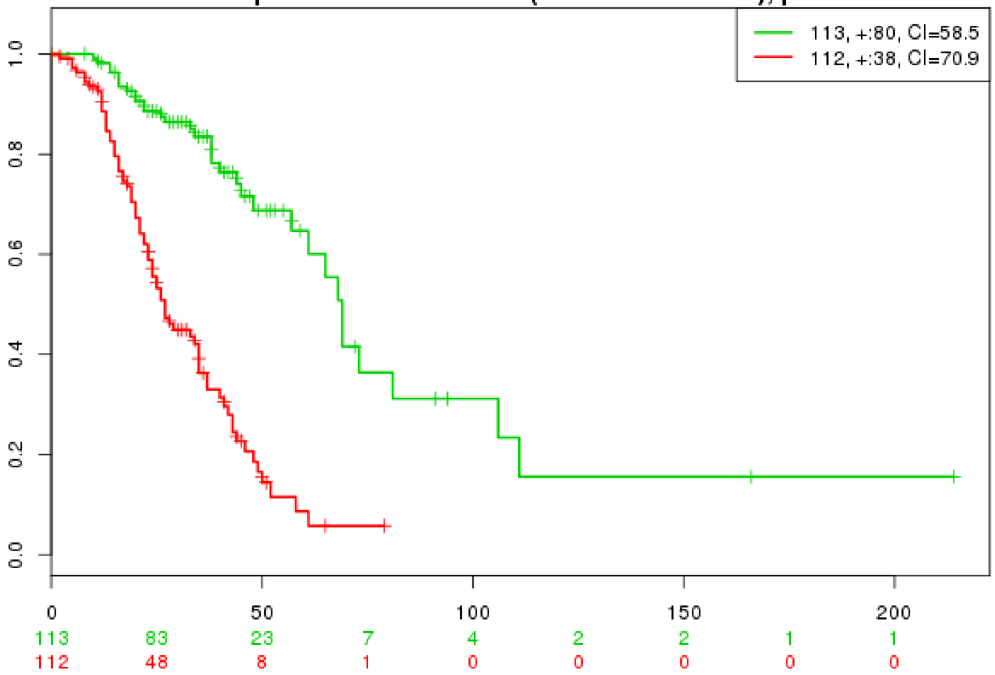

### Lightpink3 module replication (23 unique genes)

**Concordance Index = 65.7, Log-Rank Equal Curves p=7.567e-05, R^2=0.121/0.986**
**Risk Groups Hazard Ratio = 2.19 (conf. int. 1.47 ~ 3.27), p=0.0001266**

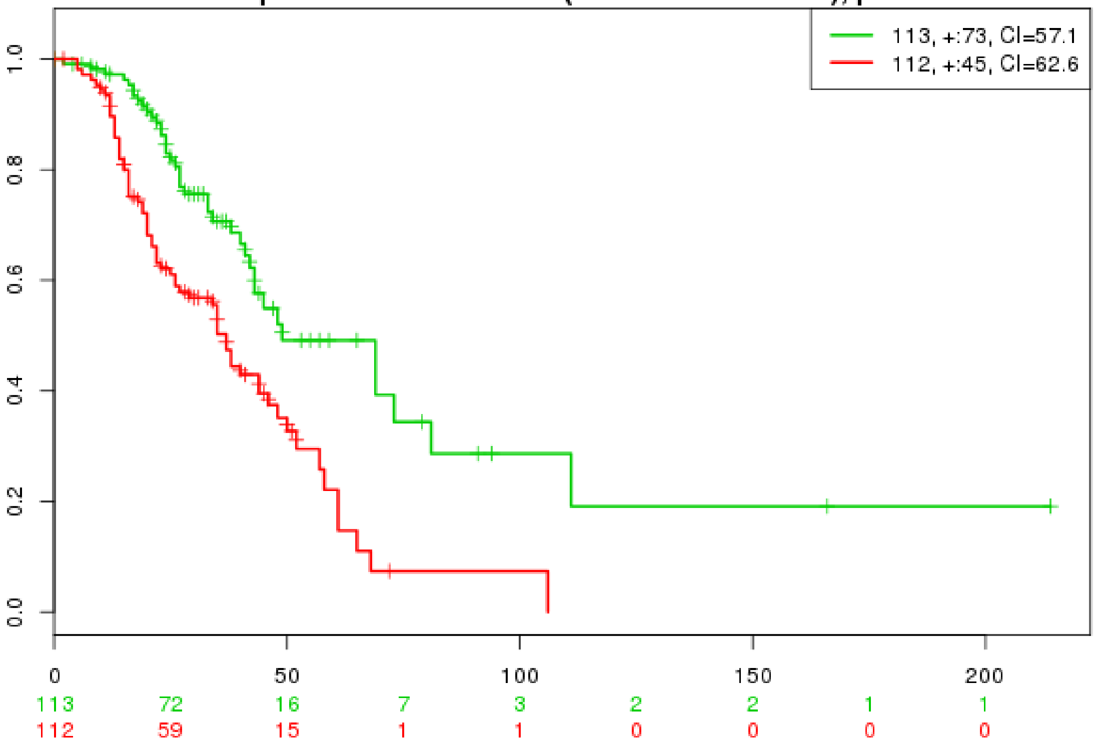


### Orangered4 module replication (34 unique genes)

**Concordance Index = 64.61, Log-Rank Equal Curves p=0.0003944, R^2=0.111/0.986**
**Risk Groups Hazard Ratio = 2 (conf. int. 1.35 ~ 2.96), p=0.0005661**

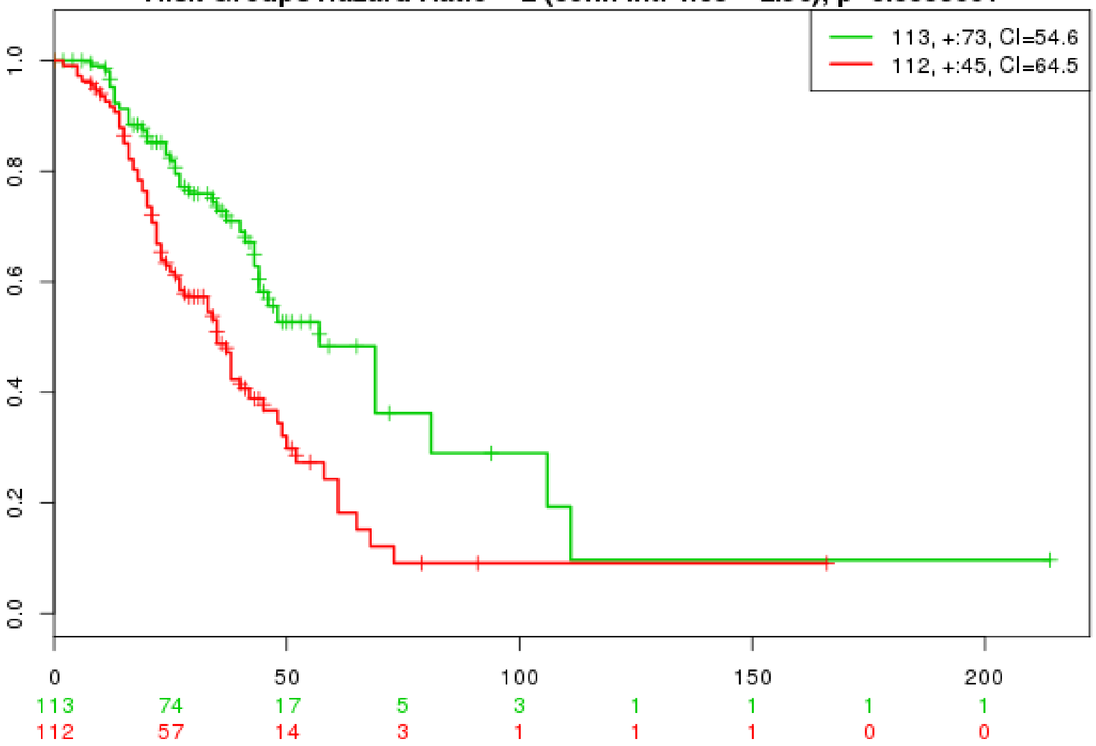

### Skyblue3 module replication (40 unique genes)

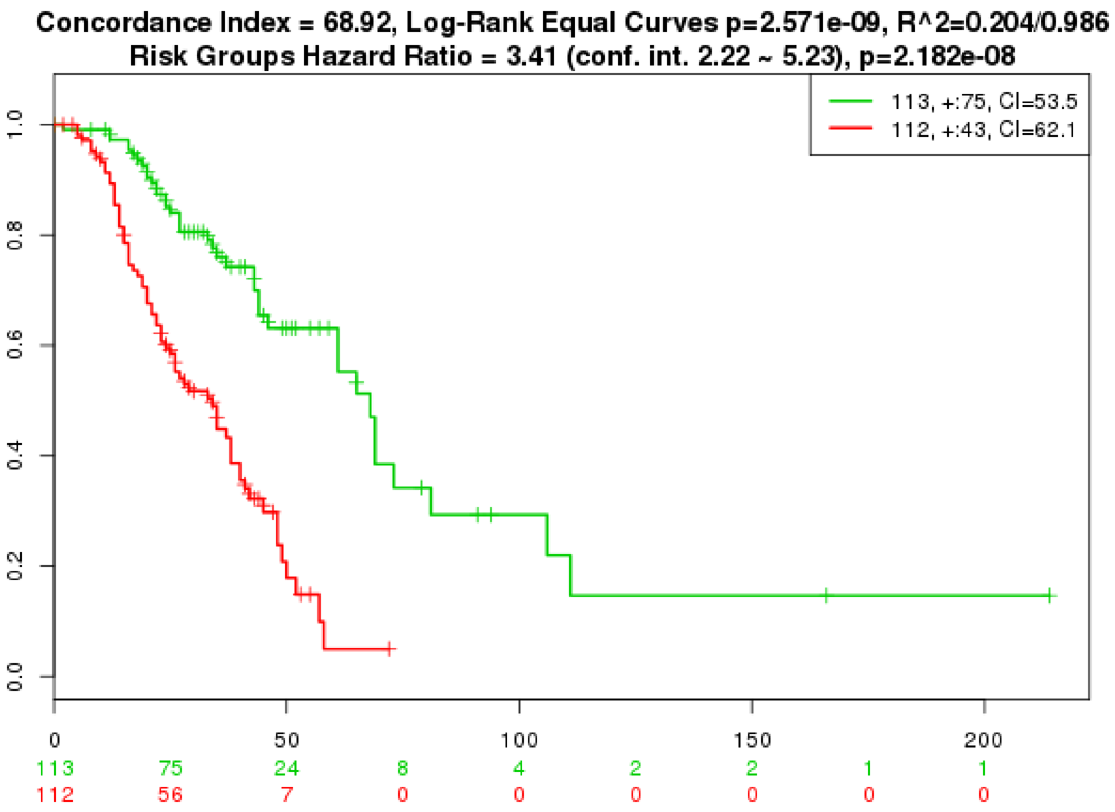


Note:

Expression of genes within each module were tested for an association with progression free survival (PFS) in an independent replication cohort (GSE9891).

Total number of study population = 225, after filtering for treatment (platinum) and histology (serous).

Replication of results were successful in all 5 significant modules, where down regulation of genes in all 5 modules are associated with poor survival rate. This is in alignment with our finding that identified co-expressed genes are downregulated in individuals who develop resistance to platinum chemotherapy.
