## Supplementary figures and images for "Gene networks and expression quantitative trait loci associated with platinum-based chemotherapy response in high-grade serous ovarian cancer"

### Supplemental Figure 5

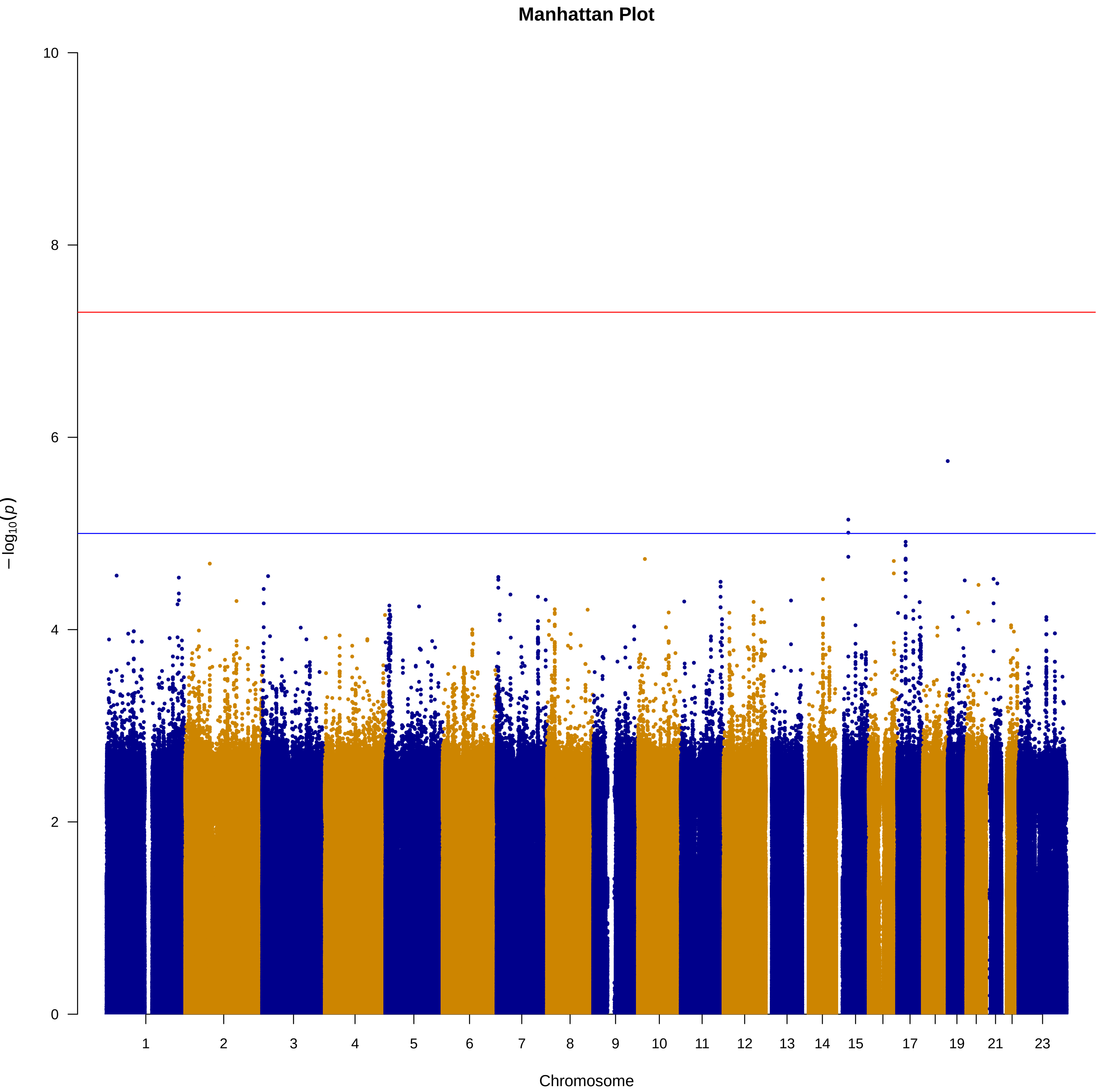
