## Supplemental Data 1 for "Gene networks and expression quantitative trait loci associated with platinum-based chemotherapy response in high-grade serous ovarian cancer"

### Microarray_Preprocessing

### 1. Loading phase

#### Loading of the required libraries

**TCGAbiolinks**

Queries and downloads relevant data from the Genomic Data Commons (GDC) of the National Cancer Institute (NCI).

**ArrayQualityMetrics**

Performs a series of comprehensive quality checks on the microarray dataset.

**Affy**

Contain functions for pre-processing Affymetrix microarray dataset.

**limma**

Used for differential expression analysis of gene expression microarray data.

```
#library(TCGAbiolinks)
#library(arrayQualityMetrics)
library(affy)
library(limma)
library(genefilter)
```

Barcodes of sensitive/resistant patients identified from the TCGA-OV cohort is loaded.

```
# read in TCGA patient barcodes
OV_sensitive <- read.table("sensitive_barcodes.txt",as.is =
TRUE)
OV_resistant <- read.table("resistant_barcodes.txt",as.is =
TRUE)
```

Using TCGABiolinks package, raw microarray data (HT_HG-U133A) from the patient’s primary solid tumor is downloaded from the NCI GDC database.

```
# search gdc & download the raw microarray data
OV_sensitive_query <- GDCquery(barcode = OV_sensitive$V1,
                          project = c("TCGA-OV"),
                          data.category = "Raw microarray
data",
```

```
                          data.type = "Raw intensities",
                          legacy = TRUE,
                          platform = c("HT_HG-U133A"),
                          sample.type = "Primary solid Tumor"
)
OV_resistant_query <- GDCquery(barcode = OV_resistant$V1,
                          project = c("TCGA-OV"),
                          data.category = "Raw microarray
data",
                          data.type = "Raw intensities",
                          legacy = TRUE,
                          platform = c("HT_HG-U133A"),
                          sample.type = "Primary solid Tumor"
)

GDCdownload(OV_sensitive_query, method = "api",directory =
"TCGA_OV_sensitive_data")
GDCdownload(OV_resistant_query, method = "api",directory =
"TCGA_OV_resistant_data")
```

### 2. Preprocessing phase

Read affy files

1. Using the Affy package, read in the affy files (.CEL) into a single affybatch variable.
2. Assign phenotype to each batch, 0 for chemo-sensitive and 1 for chemo-resistant
3. Combine the two batches for pre-processing.

```
# Read in the affy files
OV_sensitive_list <- list.files("./TCGA_OV_sensitive_data/",
".*CEL",recursive = TRUE,full.names = TRUE)
OV_resistant_list <- list.files("./TCGA_OV_resistant_data/",
".*CEL",recursive = TRUE,full.names = TRUE)

Sens_Batch <- ReadAffy(filenames = OV_sensitive_list)
Res_Batch <- ReadAffy(filenames = OV_resistant_list)
```

```
# Assigning phenotype - sensitive as 0, resistant as 1
Sens_Batch@phenoData@data$sample<-0
Res_Batch@phenoData@data$sample<-1


# Combine two phenotypes
CombinedBatch <- merge (Sens_Batch, Res_Batch)
# 553536 probes exists here (PM & MM)
```

##### Background correction & normalization

Normalization removes systematic biases and makes comparisons between arrays more meaningful. We use RMA background correction and quantile normalization.

```
#Background correction
# outputs log2-transformed expression values
# "normalize" flag - logical value. If TRUE normalize data
using quantile normalization
# "background" flag - logical value. If TRUE background
correct using RMA background correction
eset.rma = rma(CombinedBatch, background = TRUE, normalize =
TRUE)
## Background correcting
## Normalizing
## Calculating Expression
#22277 probes here


 #Control probes in the array will be removed using
featureFilter (probe starting with "AFFX_")
#Also remove probes with no Entrez Gene identifiers
eset.rma.filtered<-featureFilter(eset.rma, require.entrez=F,
remove.dupEntrez=F)
#22215 probes here
```

##### Quality check

Using the arrayQualityMetrics library, we perform a quality assessment of the array.

```
arrayQualityMetrics(eset.rma.filtered, force = TRUE, outdir =
'report')
```

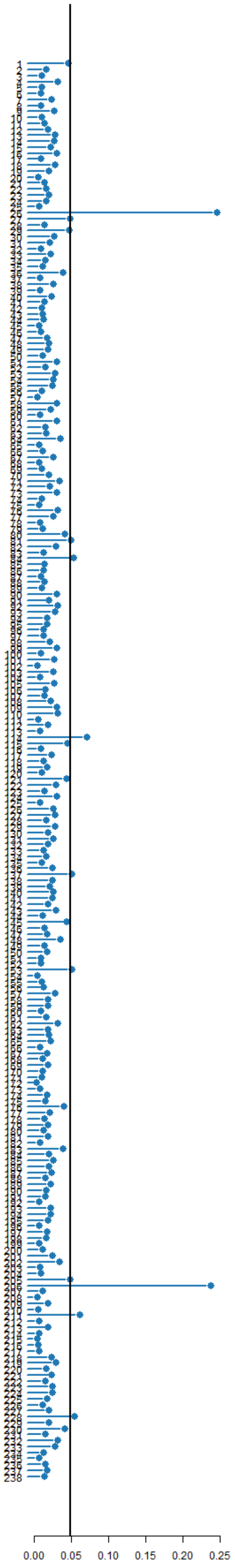
0.00
0.05
0.10
0.15
0.20
0.25

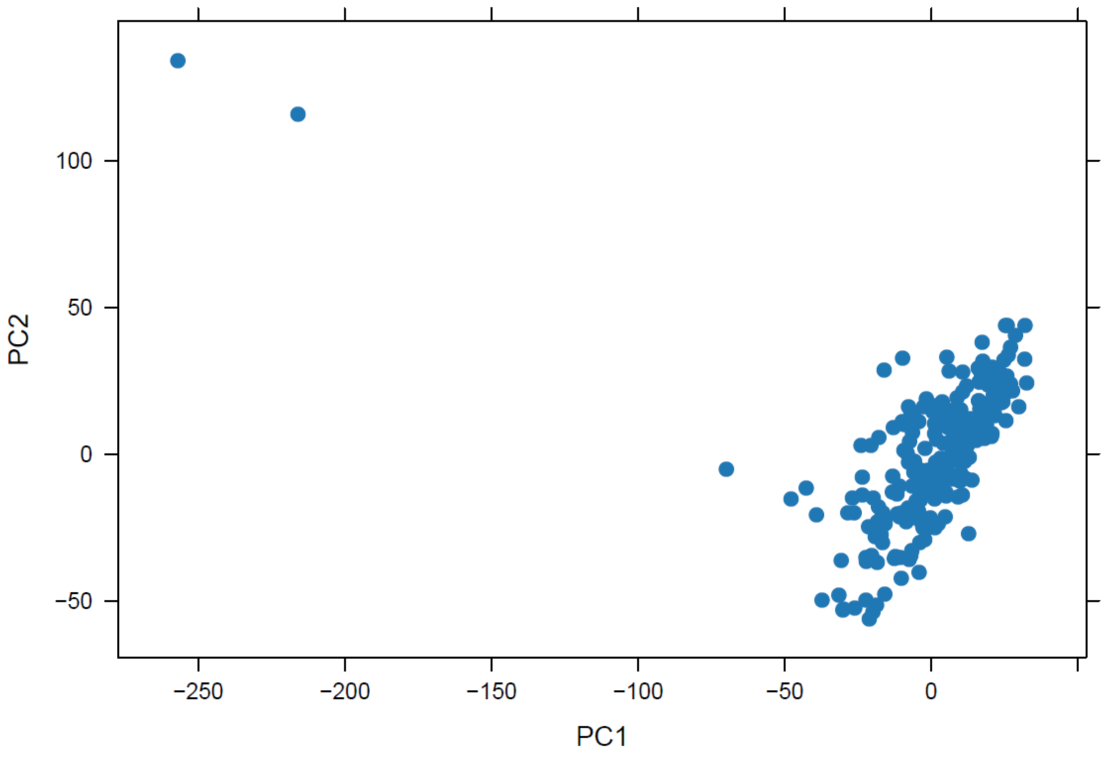

PC2
PC1
100
50
0
-50
-250
-200
-150
-100
-50
0


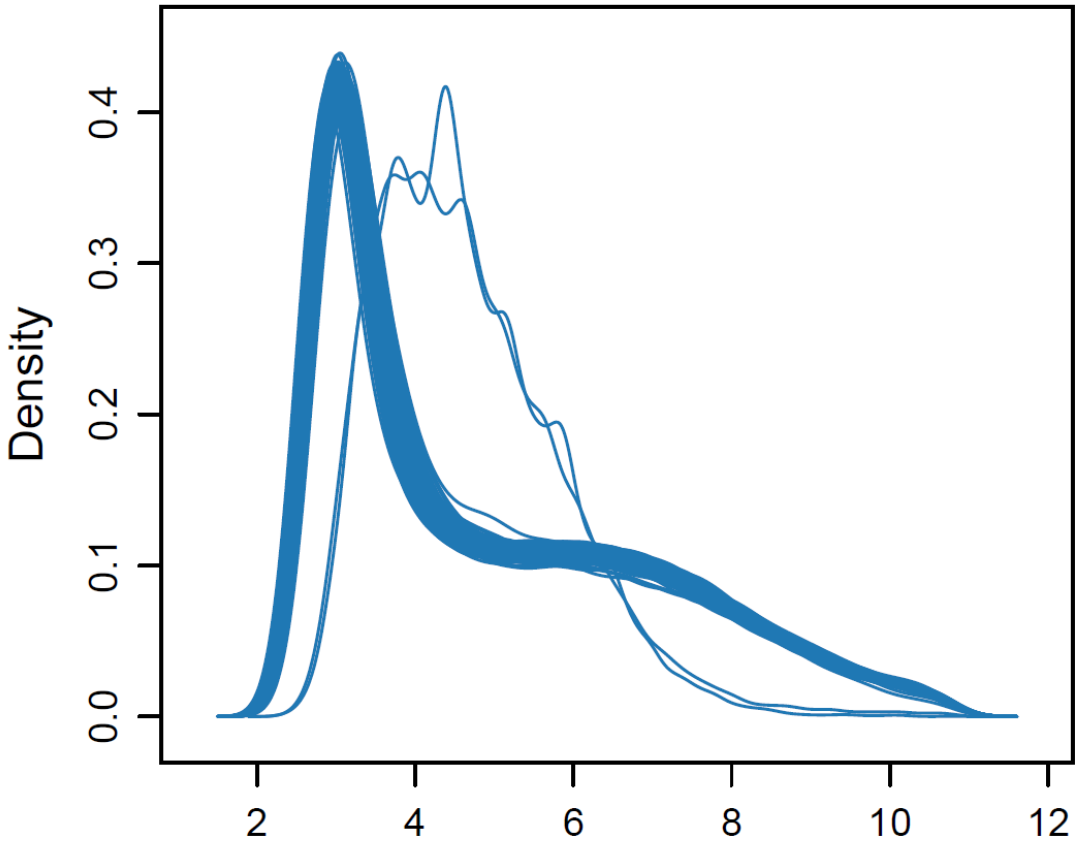

Density
0.0
0.1
0.2
0.3
0.4
2
4
6
8
10
12

From the inspection of an outlier detection, it is evident that two subjects, num.26 ("FEAST_p_TCGA_B20_21_Expression_HT_HG-U133A_96-HTA_F04_516474.CEL") and num.206 ("FEAST_p_TCGA_B20_21_Expression_HT_HG-U133A_96-HTA_G06_516372.CEL"), are strong outliers.

We therefore dropped these two subjects from our population pool.

Two duplicated samples that map to same subject were also removed.

```
# Identify the two outliers

potential_outlier1 =
match("FEAST_p_TCGA_B20_21_Expression_HT_HG-U133A_96-
HTA_F04_516474.CEL", sampleNames(eset.rma.filtered))

potential_outlier2 =
match("FEAST_p_TCGA_B20_21_Expression_HT_HG-U133A_96-
HTA_G06_516372.CEL", sampleNames(eset.rma.filtered))


# Remove two duplicate subjects we identified (from manual
check)

Duplicate1 = match("TARRE_p_MultiPlate_TCGA_SS_MA_Ref_HT_HG-
U133A_96-HTA_A06_586078.CEL", sampleNames(eset.rma.filtered))

Duplicate2 = match("AGARS_p_TCGA_B12_RNA_ReDo_HT_HG-U133A_96-
HTA_C07_443130.CEL",

sampleNames(eset.rma.filtered))


# remove from the expression set

eset.rma.outlier_controlled = eset.rma.filtered[,-
potential_outlier1]

eset.rma.outlier_controlled = eset.rma.outlier_controlled[,-
potential_outlier2]

eset.rma.outlier_controlled = eset.rma.outlier_controlled[,-
Duplicate1]

eset.rma.outlier_controlled = eset.rma.outlier_controlled[,-
Duplicate2]
```

After background correction, normalization, and outlier filtering, 234 subjects (135 sensitive and 99 resistant) and 22,215 probes remained. This expression matrix exported for further analysis.

```
dim(eset.rma.outlier_controlled) # number of ppl & probes
## Features  Samples
##    22215      234
table(eset.rma.outlier_controlled@phenoData@data$sample) # 0 = sensitive, 1 = resistant
##
##   0   1
## 135  99
```
