## Supplemental Data 2 for "Gene networks and expression quantitative trait loci associated with platinum-based chemotherapy response in high-grade serous ovarian cancer"

### Age at diagnosis

| | |
|---|---|
| Age average | 59.4 |
| Age standard deviation | 11.2 |
| Age median | 58 |
| Age min | 30 |
| Age max | 87 |

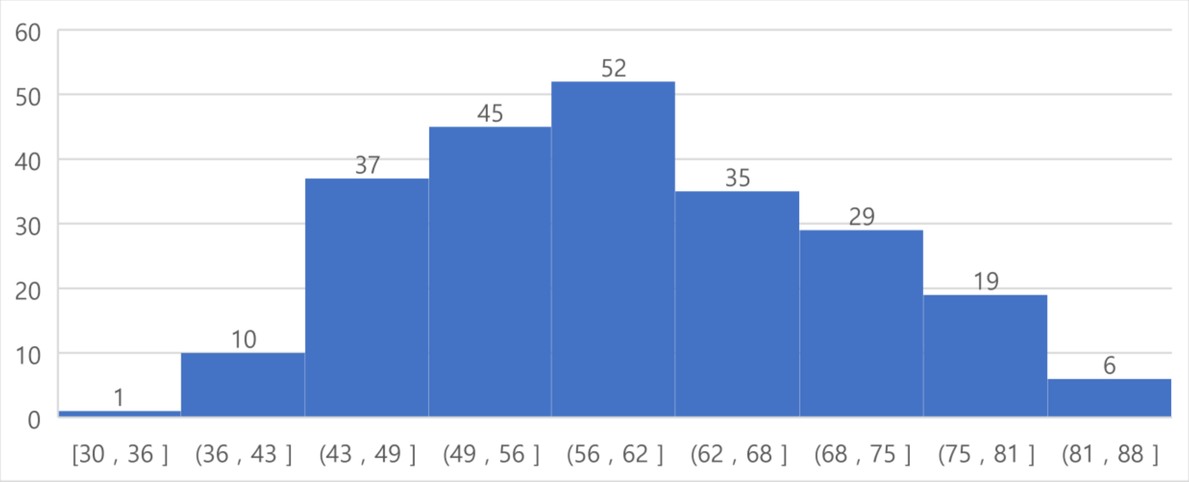

### Tumor stage at diagnosis

| Tumor Stage | Number of cases |
|---|---|
| Stage IC | 2 |
| Stage IIA | 2 |
| Stage IIC | 8 |
| Stage IIIA | 4 |
| Stage IIIB | 10 |
| Stage IIIC | 177 |
| Stage IV | 31 |
| **Total** | **234** |

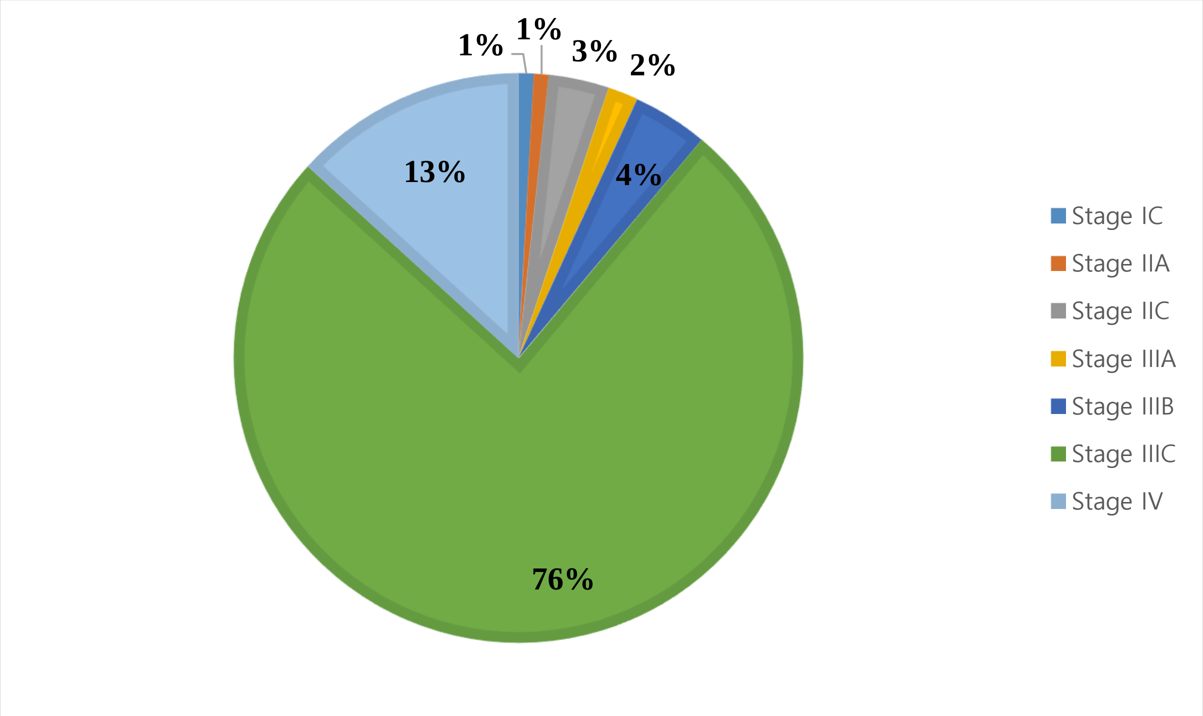
