## Supplemental Data 3 for "Gene networks and expression quantitative trait loci associated with platinum-based chemotherapy response in high-grade serous ovarian cancer"

### TCGA-OV genotype imputation log

#### Variant imputation

Michigan imputation server (https://imputationserver.sph.umich.edu) was used to impute for missing genotypes.

##### Autosomes (chr 1 – 22)

Chromosomes: 1 2 3 4 5 6 7 8 9 10 11 12 13 14 15 16 17 18 19 20 21 22
SNPs: 868263
Chunks: 153
Datatype: unphased
Reference Panel: phase3
Phasing: eagle

**Statistics:**
Alternative allele frequency > 0.5 sites: 235,699
Reference Overlap: 99.67%
Match: 860,761
Allele switch: 0
Strand flip: 0
Strand flip and allele switch: 0
A/T, C/G genotypes: 0
**Filtered sites:**
Filter flag set: 0
Invalid alleles: 448
Duplicated sites: 2
NonSNP sites: 0
Monomorphic sites: 1,088
Allele mismatch: 3,105
SNPs call rate < 90%: 0

**X chromosome**

Chromosomes: X
SNPs: 36764
Chunks: 8
Datatype: unphased
Reference Panel: phase3
Phasing: shapeit

**Statistics:**
Alternative allele frequency > 0.5 sites: 0
Reference Overlap: 99.05%
Match: 26,353
Allele switch: 7,769
Strand flip: 0
Strand flip and allele switch: 0
A/T, C/G genotypes: 1,383
**Filtered sites:**
Filter flag set: 0
Invalid alleles: 0
Duplicated sites: 0
NonSNP sites: 0
Monomorphic sites: 785
Allele mismatch: 133
SNPs call rate < 90%: 0

**Total number of variants**

| | **Autosomes** | **X-chromosome** |
|---|---:|---:|
| **Before imputation** | 868,263 | 36,764 |
| **After imputation** | 46,181,723 | 1,778,607 |
