## Supplemental Data 4 for "Gene networks and expression quantitative trait loci associated with platinum-based chemotherapy response in high-grade serous ovarian cancer"

### TCGA-OV genotype data QC pipeline

Plink (v. 1.9) was used to perform quality check of the genomics dataset.

Sample level:

- Genetic sex was determined based on heterozygosity rates of SNPs on sex chromosomes. In total, four samples were marked as sex outliers (F > 0.2) and not females, which were removed from the study.

- Two samples with genotyping call rate less than 90% were removed.

- Genetic relatedness was calculated and two pairs of highly related individuals were identified (pi-hat > 0.9). One individual from each pair was randomly selected and removed.

- Observed and expected autosomal homozygous genotype counts were computed for each subject to obtain a F coefficient estimate. A total of 20 samples with either high (F> 0.05) or low (F< -0.05) heterozygosity rates were excluded.

**These steps removed 26 subjects in total. Thus, of the 266 subjects (157 sensitive and 109 resistant) with available genotype dta, 240 subjects (142 sensitive and 98 resistant) passed QC measures.**

Variant level:

- 772 variants with more than 10% missingness in genotyping rate were removed.

- 38,430,595 rare variants with minor allele frequency less than 1% were excluded from analysis.

**These steps removed 8,431,367 variants in total. Thus, of the 47,960,330 variants (after imputation), 9,528,963 SNPs passed QC measures.**
